# The 100,000 most influential scientists rank: the underrepresentation of Brazilian women in academia

**DOI:** 10.1101/2020.12.22.423872

**Authors:** Leticia de Oliveira, Fernanda Reichert, Eugenia Zandona, Rossana C. Soletti, Fernanda Staniscuaski

## Abstract

Despite the progress observed in recent years, women are still underrepresented in science worldwide, especially at top positions. Many factors contribute to women progressively leaving academia at different stages of their career, including motherhood, harassment and conscious and unconscious discrimination. Implicit bias plays a major negative role in recognition, promotions and career advancement of female scientists. Recently, a rank on the most influential scientists in the world was created based on several metrics, including the number of published papers and citations. Here, we analyzed the representation of Brazilian scientists in this rank, focusing on gender. Female Brazilian scientists are greatly underrepresented in the rank (11% in the Top 100,000; 18% in the Top 2%). Male scientists have more self-citation than female scientists and positions in the rank varied when self-citations were included, suggesting that self-citation by male scientists increases their visibility. Moreover, male scientists had more papers never cited than female scientists. Possible reasons for this observed scenario are related to the metrics used to rank scientists, since these metrics reproduce and amplify the well-known implicit bias in peer-review and citations. Discussions on the repercussions of such ranks are pivotal to avoid deepening the gender gap in science.

## INTRODUCTION

In Brazil, women are deeply underrepresented at the senior and leadership levels in academia, especially when considering decision-making positions (Valentova et al. 2017; Areas et al. 2020). The percentage of women decreases disproportionately as they progress in their careers, which is a globally-observed phenomenon (Frietsch et al. 2009) and known as vertical or hierarchical segregation (Rossiter 1982), scissors effect (van Vlooten 2005; Areas et al. 2020) or leaky pipeline (Pell 1996). Etzkowitz & Ranga (2011) state that the leaky pipeline “emphasises a linear progression through a series of staged roles within academia, with a loss of female talent at every critical transition”. There are several factors that contribute to women progressively leaving academia throughout their career, such as motherhood, domestic labor, child care (Machado et al. 2019; Frietsch et al. 2009), harassment (National Academies of Sciences, Engineering, and Medicine 2018) and conscious and unconscious gender bias among others (Moss-Racusin et al. 2012; Reuben et al. 2014; Gaston 2015; Carli et al. 2016). Implicit bias against women, which is an unconscious belief that women are less capable than their male peers, causes considerable damage to the progress of their scientific careers (Moss-Racusin et al. 2012; Dutt et al. 2016; Kuo 2016). For instance, experimental studies have shown that CVs with a male name are evaluated as more competent and deserving a higher salary than the same CV with a female name (Moss-Racusin et al. 2012; Eaton et al. 2020).

A traditional way to measure productivity and prestige in academic science is through publications and citations (Murray & Graham 2007), which are used to evaluate scientists for hiring, promotion, and funding (West et al. 2013). As an example, among first-time Principal Investigators awarded with all types of National Institutes of Health (NIH) grants from 2006 to 2017, women received a median of $126,615 vs $165,721 for men (Oliveira et al. 2019). These funding disparities may compromise women’s future research performance, turning them less competitive and thus harming their persistence in academia. High-impact journals, such as those from the Nature group, have much less women as senior authors (18.1% in last authorship), proportion that decreases with increasing impact factor of the journal (Bendels et al. 2018). Importantly, however, when articles are reviewed anonymously (double-blind review), the number of articles published with women as first authors increases (Budden et al. 2008). In addition, articles with women as leading authors are less cited than those with men as leading author (Larivière et al. 2013; Dworkin et al. 2020; Elsevier 2020). All of these examples highlight how implicit bias can negatively impact the publication and citation processes.

Recently, a rank identifying the 100,000 most influential scientists in the world was published (Ioannidis et al. 2019; 2020). This list can have a great impact on the career of scientists, as such visibility can have implications for networking and for obtaining research funding. The authors used Scopus data to identify a database of the 100,000 most cited authors in all scientific areas based on a composite indicator that considers six citation metrics: (1) total citations; (2) Hirsch h-index; (3) coauthorship-adjusted Schreiber hm-index; (4) number of citations of single-author papers; (5) number of citations of single-author or first-author papers; and (6) number of citations of single-author, first-author, or last-author papers (Ioannidis et al. 2016). They provided ranks with and without self-citations. Moreover, they presented a rank that considers the Top 2% scientists of their main subfield disciplines. Ranks are presented for career-long and single-year impact. Here, we analyzed the representation of Brazilian scientists by gender in the database built by Ioannidis et al. (2020) to investigate a possible underrepresentation of women scientists. Considering that women are typically underrepresented in academia, especially in higher-rank positions (van den Besselaar & Sandström 2017), we expect that Brazilian women will be greatly underrepresented in the 100,000 top scientists recently-published ranking. It is our aim to raise awareness to the problem of a gender-biased scientific elite and to the validity of the metrics that have been typically used in academia.

## METHODS

The study was based on the publicly available database of 100,000 top scientists developed by Ioannidis and collaborators (2019; 2020). The database was downloaded from https://data.mendeley.com/datasets/btchxktzyw/2 (Bass et al. 2020). This database shows the 5 domain, 20 field and 174 subfields of scientists and presents two independent ranks: (1) Top 100,000 rank and (2) a rank that considers the Top 2% scientists of their main subfield disciplines. Ranks are described including or excluding self-citations, and are presented considering the career-long (Career dataset) or single-year impact (2019). The Career dataset considers publications and citations from 1960 to 2019, while the single-year dataset takes into consideration the metrics only for 2019. As our aim was to analyze the representation of Brazilian scientists by gender, all datasets were first filtered by “Country”, in order to select only those that are currently affiliated to Brazilian institutions. Since there is no explicit information on gender in the datasets, gender was attributed to each of the scientists in the list based on their first names. Ideally, an inclusive gender classification system should be used. Unfortunately, based on information available in the database used in this study, it was only possible to use the binary classification (male or female), since first names are embedded into the gender binary. This is a limitation, but as long as datasets lack gender identification, name-based gender inference remains the method of choice for plenty of applications, including studies of women’s representation in science (Santamaria & Mihajjevic 2018). We manually labeled the first names as male or female according to Brazilian culture, where names are commonly gender-specific. When only the initials were available or in case of uncommon first names, we performed an internet search (i.e., Google Scholar, Curriculum Lattes platform, Research Gate, LinkedIn, universities websites, etc.) to confirm the gender identity of the researcher, crossing multiple information available on the database (last name, initials, affiliation institution and research field). We could not assign gender for only one of the Brazilian researchers listed in the database, due to the impossibility to find out the researcher’s first name. Three Brazilian scientists were duplicated in the Single Year dataset. For the analysis performed here, the duplicated entries were excluded. To analyze the representation of Brazilian scientists in the two ranks (Top 100,000 rank and the rank that considers the Top 2% scientists of their main subfield disciplines), we described the total number of Brazilian scientists, as well as the number of male and female scientists, in each one. Moreover, the data were analyzed considering career-long (Career dataset) and single-year impact (2019). We also analyzed separately the data that included or excluded self-citations. For the subsequent analysis, where we evaluated self-citation percentages, number of published papers, number of never-cited published papers and the scientific field, we combined all Brazilian scientists into one single list and analyzed them regardless of each rank they were originally from. The number of published papers, self-citation percentage and scientific domain, field and subfield for each scientist are provided in the original database. The number of never-cited papers was calculated as the difference between the number of published papers and the number of cited papers from 1960 to 2019 that is provided in the database. To evaluate the impact of self-citation in rank position, we used the information of rank position present in the database for both conditions (including and excluding self-citations) and calculated the percentage of male and female scientists that increased, decreased or did not change their position in the rank.

A student t-test was performed to evaluate statistical differences between men and women in the average self-citation index, the average number of papers published and the average number of papers never cited, both including and excluding self-citations. The significance level was set at 0.05. All analyses were performed using SigmaPlot version 10.0, from Systat Software.

## RESULTS

### Representation of Brazilian scientists in the ranks

We have analyzed all the available datasets from the database in order to determine Brazil’s participation in the ranks. In the Top 100,000 rank, excluding self-citations, there are 254 Brazilian scientists in the Career dataset and 352 in the Single Year dataset, representing 0.25% and 0.35%, respectively, of the world’s top 100,000 scientists. When self-citations are included, the participation of Brazilian scientists increases to 0.3% (Career dataset, 302 scientists) and 0.39% (Single year dataset, 391 scientists). In the rank that considers the Top 2% scientists of their main subfield disciplines, Brazilian researchers correspond to 0.38% (Career dataset, 600 scientists) and 0.53% (Single year dataset, 853 scientists) of the world’s most influential scientists (Figure 1).

**Figure 1.**
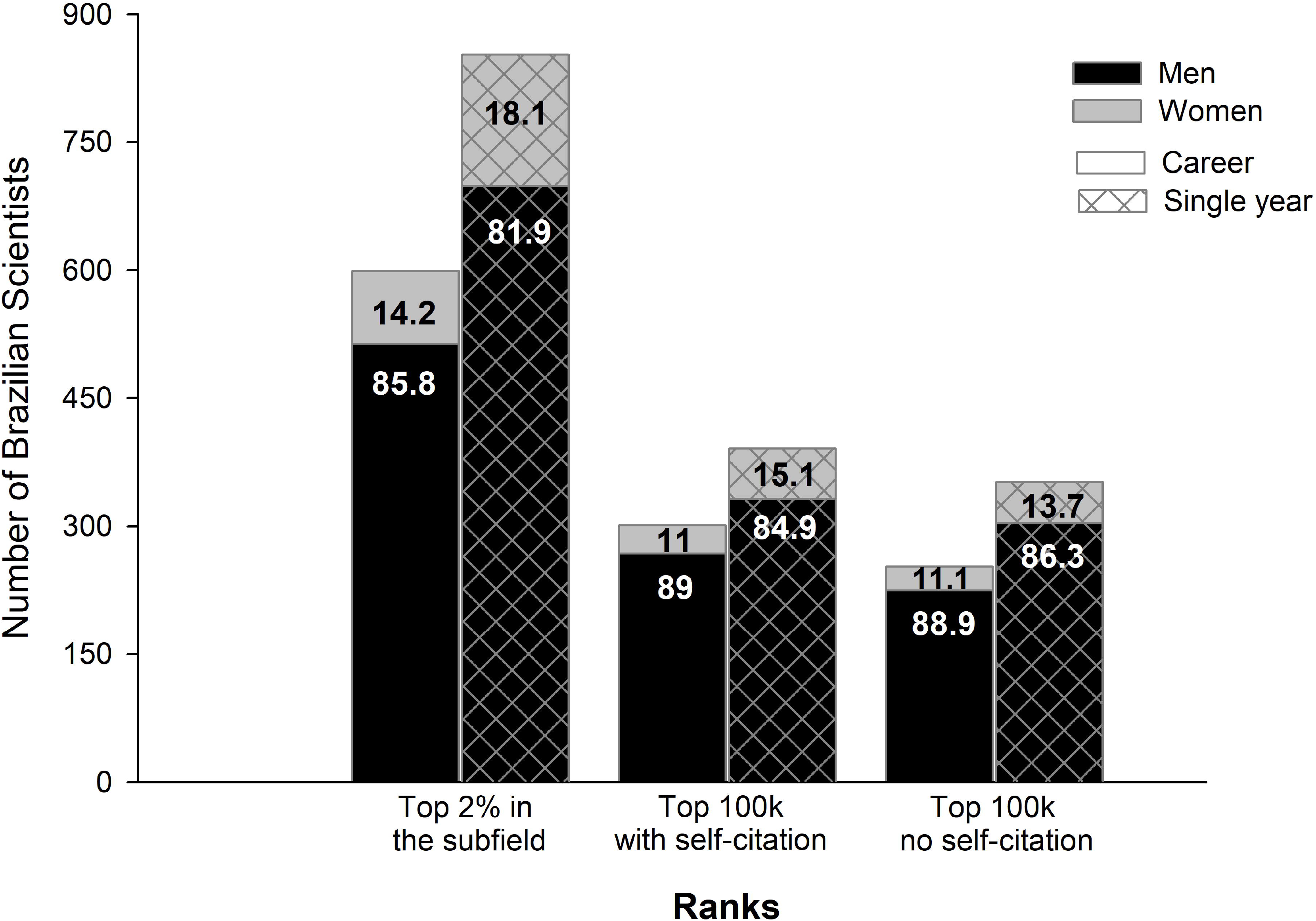
Representation of Brazilian researchers by gender in the Most Influential Scientists Rank. Number of Brazilian Scientists by gender in each of the ranks. Numbers in the bars indicate the percentage of male and female scientists.

When analyzing data by gender, we observed a great disparity between men and women (Figure 1). In both Top 100,000 ranks, including and excluding self-citations, women represent only 11% of the Brazilian scientists in the Career dataset. In the Single Year dataset, women represent 15.1% of Brazilian researchers in the rank including self-citations, and 13.7% in the rank excluding self-citations. Although still greatly underrepresented, percentages of women are higher in the Top 2% rank, being 14.2% and 18.1% of Brazilian scientists in the Career and Single Year datasets, respectively.

### Representation of Brazilian scientists by field

In total, regardless of the ranks they were originally from, there are 1022 Brazilian scientists in the database presented by Ioannidis et al. (2020). These scientists are distributed in all five scientific domains, but there is a great imbalance in the representation of each domain. The domains with the highest numbers of Brazilian scientists are Health Sciences, Natural Sciences, and Applied Sciences (with 384, 356 and 272 Brazilians respectively). The domains with the lowest representation of Brazilian scientists are Economic & Social Sciences with eight, and Arts & Humanities with only two. Considering gender, the highest percentage of Brazilian women was found in Health Sciences (21.9% of Brazilian scientists). This percentage decreases to 17.3, 13.8 and 12.5% for Applied Sciences, Natural Sciences, and Economic & Social Sciences, respectively. There are no Brazilian women in the Arts & Humanities domain. Brazilian scientists are represented in 18 of the 20 fields, with no representation in Communication & Textual Studies and Visual & Performing Arts. The fields with the highest number of Brazilians are Clinical Medicine, Chemistry, Physics & Astronomy, and Biology (Figure 2). Considering gender, in 30% of the fields there are no Brazilian women, and there is no field with only female scientists. For the fields in which both male and female scientists are represented, the highest number of women is observed in Agriculture, Fisheries & Forestry (27.5%) and Public Health & Health Services (25%), while the lowest are in Physics & Astronomy (5.8%) and Engineering (8.1%).

**Figure 2.**
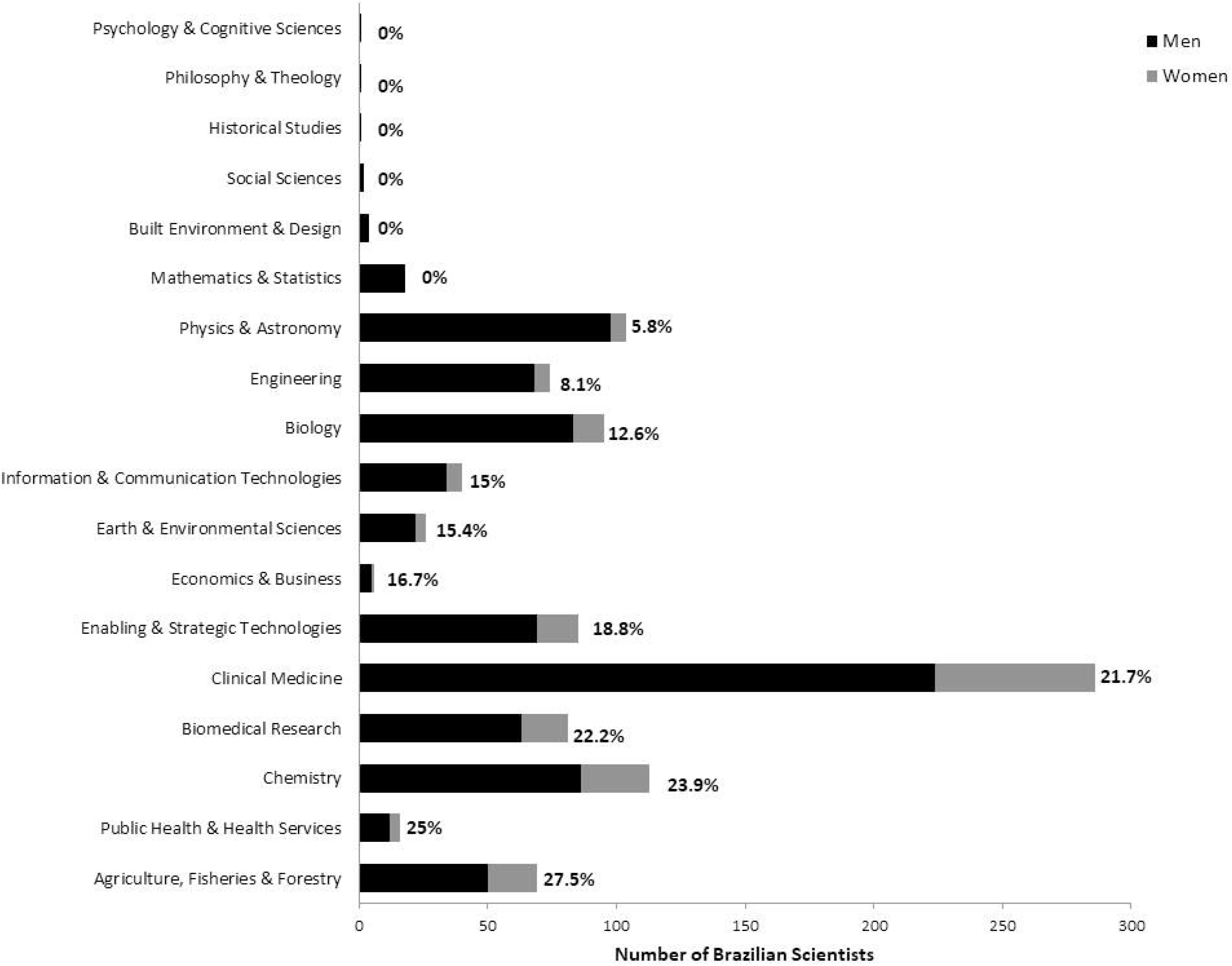
Representation of Brazilian researchers in the ranks by field and gender. Number of Brazilian Scientists in each of the scientific fields by gender. Numbers next to the bars indicate the percentage of female scientists. The two fields (Communication & Textual Studies and Visual & Performing Arts) in which there were no Brazilian scientists are not shown.

Considering the subfield classification, Brazilian scientists appear in 106 subfields, with the overall highest numbers in Dentistry (60 researchers), Tropical Medicine (49 researchers) and Medical & Biomolecular Chemistry (42 researchers) (Supplementary figure 1). Brazilian women are not present in 51 (48%) of the subfields. In contrast, only two subfields (1.9%) - Nanoscience & Nanotechnology and Anesthesiology - are only represented by Brazilian women. In the subfields where both men and women are represented, the highest numbers of women are in Nutrition & Dietetics (75%) and General Clinical Medicine (66.7%). In contrast, the lowest numbers of Brazilian women are observed for Nuclear & Particle Physics (3.6%) and Materials (4.5%) subfields.

### Self-citation influence on Brazilian scientists impact

Several factors influenced the inclusion of scientists in the ranks, including self-citations percentage. In general, Brazilians present a higher self-citation percentage (19.3%) than the world’s average (13.7%). When factoring gender, there is a significant difference (Student t-test, p = 0.008) between self-citations percentage of female (18%) and male (19.6%) Brazilians (Figure 3a). Positions in the rank varied greatly when comparing the datasets including or excluding self-citations (Figure 3b). Self-citation inclusion moved 67.7% and 61.9% of Brazilian male and female scientists up in the ranks, respectively, while 32 and 38.1% of Brazilian male and female scientists moved down in the ranks when self-citations were considered. On average, Brazilian scientists published less papers (173) in the period (1960 - 2019) when compared to the world’s average (198). A significant difference between the number of published papers was observed between male (180) and female (141) Brazilian scientists (Student t-test, p = 0.00005; Figure 3c). It was interesting to note that the percentage of papers that were never cited was significantly higher for men (21.6%) than that for women (19.1%) when self-citations were not considered (Student t-test, p = 0.02; Figure 3d). If self-citations are considered, there is no longer a significant difference between the percentage of never-cited papers for male (17%) and female (15.4%) Brazilian scientists (Student t-test, p = 0.11).

**Figure 3.**
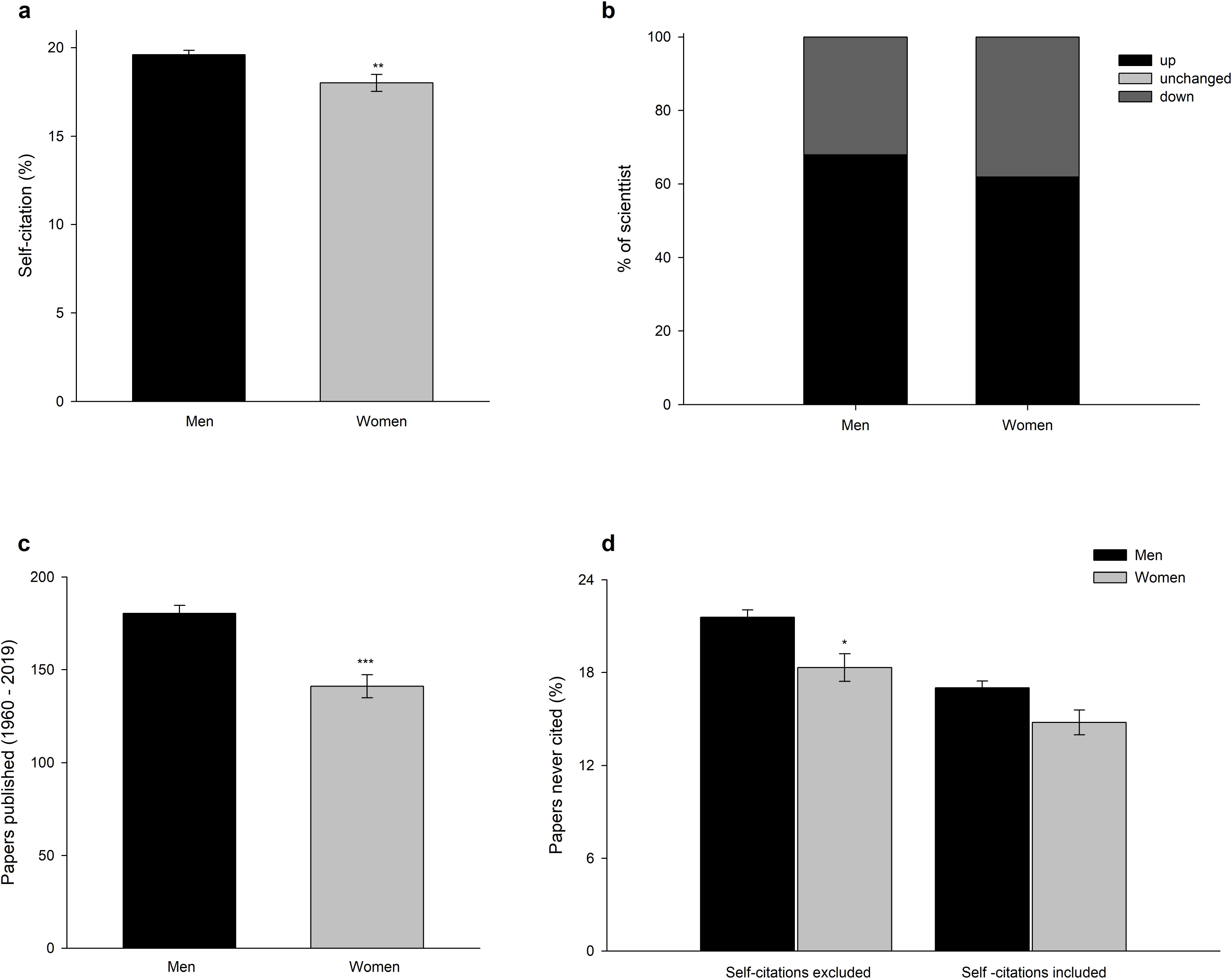
Number of published papers and self-citation impact by gender. **a**. Percentage of self-citations by Brazilian scientists by gender. **b**. Rank position changes after the inclusion of self-citations. (c) Number of papers published between 1960 and 2019. (d) Number of papers published between 1960 and 2019 that were never cited, including and excluding self-citations. Data are shown as percentage (b) or mean + standard error (a, c, d). Statistical differences in means were analyzed using Student t-tests (* p < 0.05, ** p < 0.01, *** p < 0.001).

## DISCUSSION

In the present study we investigated the representation of Brazilian scientists in the recent rank of the 100,000 most influential scientists in the world (Ioannidis et al. 2020) mostly focusing on a gender perspective. We found that Brazilians represent a very small percentage of scientists listed in the database (the highest percentage being only 0.53% for the Single year dataset). When analyzing the datasets by gender, as expected, we found that women are greatly underrepresented. Among Brazilian scientists figuring the ranks, women account for as low as 11% (Top 100,000 rank - Career dataset) and no higher than 18% (Top 2% rank - Single Year dataset). Brazilian women are underrepresented in all areas and are completely absent in some fields and subfields, even some that typically show high percentages of women, such as those related to Arts & Humanities.

We found a correspondence in the representation of Ioannidis et al. (2020) dataset and the literature. Brazil contributes with highly relevant research in several fields, such as in Medicine, Nursing, Physics & Astronomy and Dentistry (McManus et al. 2020). These fields are indeed those with the highest number of Brazilian scientists in the ranks. On the other hand, the impact of Brazilian research is lower for Social and Human Sciences (McManus et al. 2020), which is confirmed by the low representation of Brazilians for these fields in the Ioannidis et al. (2020) ranks. A recent analysis by Santiago et al. (2020) showed big disparities in the quantity of women holding a PhD between areas in Brazil, with a predominance in the Humanities (19.8% of women PhD), and a very small presence in Engineering (4.3%). In contrast, the number of female Brazilian scientists present in the ranks analyzed here does not seem to reflect this pattern, since the lowest representation observed was in the Economic & Social Science and Arts & Humanities domains. However, the presence of Brazilian scientists in these two domains is particularly low, which is a result that merits a more thorough investigation.

Our results also corroborate with findings in the literature that women are underrepresented in science, especially in top positions. This phenomenon has been previously reported as a scissors effect or leaky pipeline in the scientific career of women (van Vlooten 2005; Frietsch et al. 2009; Etzkowitz & Ranga, 2011). Specific to the Brazilian scenario, women are the majority when entering the scientific career (women represent 53% of graduate students), however, their participation decreases greatly as the career progresses (i.e.: women represent 36% research fellowship grants recipients from the National Council for Scientific and Technological Development - CNPq, 16% of the Brazilian Society for the Progress of Science - SBPC presidents and 0% of the Brazilian Academy of Sciences - ABC or CNPq presidents) (Areas et al. 2020). Possible explanations for the scissor effect in science are several and not yet fully understood. Differences in family duties (Jolly et al. 2014), in funding (Pohlhaus et al. 2011), in networking (Uhly et al. 2015), and the negative effect of implicit bias and gender role stereotypes (Moss-Racusin et al. 2012; Dutt et al. 2016; Kuo 2016) are all factors that could perpetuate women underrepresentation.

The underrepresentation of Brazilian women among the top 100,000 scientists ranking can contribute to reducing their visibility, especially for those in leadership positions. This can potentially create a vicious circle, where a lower perceived academic performance leads to less visibility, in turn making it even more difficult to increase productivity. Specifically, gender negatively affects academic position, which has a negative effect on researchers’ performance, thus, in turn, strengthening the lower status (van den Besselaar & Sandstrom 2017). This pattern could be reinforced when using productivity metrics that have a gender bias. For instance, women publish less as first and last authors and are less cited (Bendels et al. 2018; Larivière et al. 2013; Dworkin et al. 2020; Elsevier 2020), thus using metrics that are just based on number of publications and citations reproduces and reinforces the observed gender disparity in academia. Therefore, as the metrics used by Ioannidis et al. (2019; 2020) have a gender bias, by construction, the list of the most influential scientists will be composed in its majority by male scientists. Using this type of metric exclusively, only reinforces the gender bias in science.

When we analyzed the effect of self-citation with a gender perspective, we found that male Brazilian scientists in the rank have a significantly higher self-citation index than their women peers. This was also observed in an analysis of 1.5 million papers published between 1779 and 2011, where men were found to cite their own papers much more frequently than women (King et al. 2017). Moreover, our results showed that self-citations affected the position of scientists in the rank, where self-citation increases the visibility of male scientists. Furthermore, Brazilian male scientists produce more papers never cited, especially when self-citations were not included. The percentage of uncited papers of Brazilian men is comparable to a global trend - 39 million research papers across all disciplines recorded in Web of Science from 1900 to the end of 2015 - where around 21% of papers have no citations (van Noorden 2017).

However, even when women publication metrics (number of papers published and number of citations) are similar to those of their men peers, they are still less likely to become research leaders (Van Dijk et al. 2014). An analysis of almost 24,000 applications submitted to the Canadian Institutes of Health Research (CIHR) showed that when applications were evaluated primarily based on the quality of the science, the predicted probability of success was 0.9 percentage points lower for female than male applicants. However, when the evaluations were based primarily on the PI leadership and expertise, the gender gap increased to 4 percentage points (Witteman et al. 2019). Importantly, when evaluation committees of funding agencies are aware of gender bias against women, the unequal distribution of funding between men and women is less likely to occur (Régner et al. 2019). Recently, Huang et al. (2020) suggested that the gender differences observed in productivity and research impact are explained by different publishing career lengths and dropout rates. They found that the gradual increase in the presence of women in STEM over the past 60 years was paradoxically accompanied by an increased gender difference in productivity and impact, particularly among the highly productive authors. Women and men scientists publish a comparable number of papers per year and have equivalent career-wise impact for the same amount of work. The differences, however, were found in gender-specific dropout rates and the subsequent gender gaps in publishing career length and total productivity. They suggested that the community should strive to prevent the loss of women at all stages of their careers, not just junior scientists. All these examples show that implicit bias and other factors, sometimes not identified, are greatly impacting the career of women scientists, creating a series of obstacles for their permanence in academia.

In summary, we found that female Brazilian scientists are greatly underrepresented in the ranks described by Ioannidis et al. (2020) dataset. It is important to highlight that we are not questioning the merit of those Brazilian researchers who are listed in the top 100,000 scientists rank. Brazil has been suffering an ever increasing lack of funding for science and research in general and, in this context, having their international impact recognized certainly reflects the excellence of these scientists. Nevertheless, it is necessary to discuss how factors beyond merit and excellence drive the exclusion of women from such ranks. Considering the arguments presented here, we suggest that rankings of top scientists, such as that of Ioannidis et al (2020), should be published by gender. In this way, prominent women scientists could be “unmasked” by giving visibility to their scientific contributions. We believe this would help to break the cycle women scientists are facing: “less visibility - less publication and citation - less financial grants - less visibility”. If gender is not considered an important factor in the analysis of these ranks, we are deepening gender disparities in science, which is no longer acceptable. It is time to make science a fairer environment for the present and future generation.

## Acknowledgments

We would like to thank our children who are the reason we keep on the fight for a fairer world. FR acknowledges the financial support of the Brazilian National Council for Scientific and Technological Development - CNPq under the grant number 436344/2018-1. LO and EZ were supported by grants from CNPq, CAPES and FAPERJ. EZ was also supported by Prociência UERJ.

## Author Contributions

LO, EZ, FR, RCS, FS: Conceptualization, Data curation, Formal Analysis, Investigation, Methodology, Visualization, Writing – original draft, Writing – review & editing.

---

Supplementary Figure 1. **Gender distribution among Brazilian Scientists in the subfields**. Data is presented as the number of men and women in each of the subfields. Numbers next to the bars indicate the percentage of female scientists. Subfields in which there were no Brazilian scientists are not shown.

