## Supplementary figures and images for "The 100,000 most influential scientists rank: the underrepresentation of Brazilian women in academia"

### Supplementary figure 1

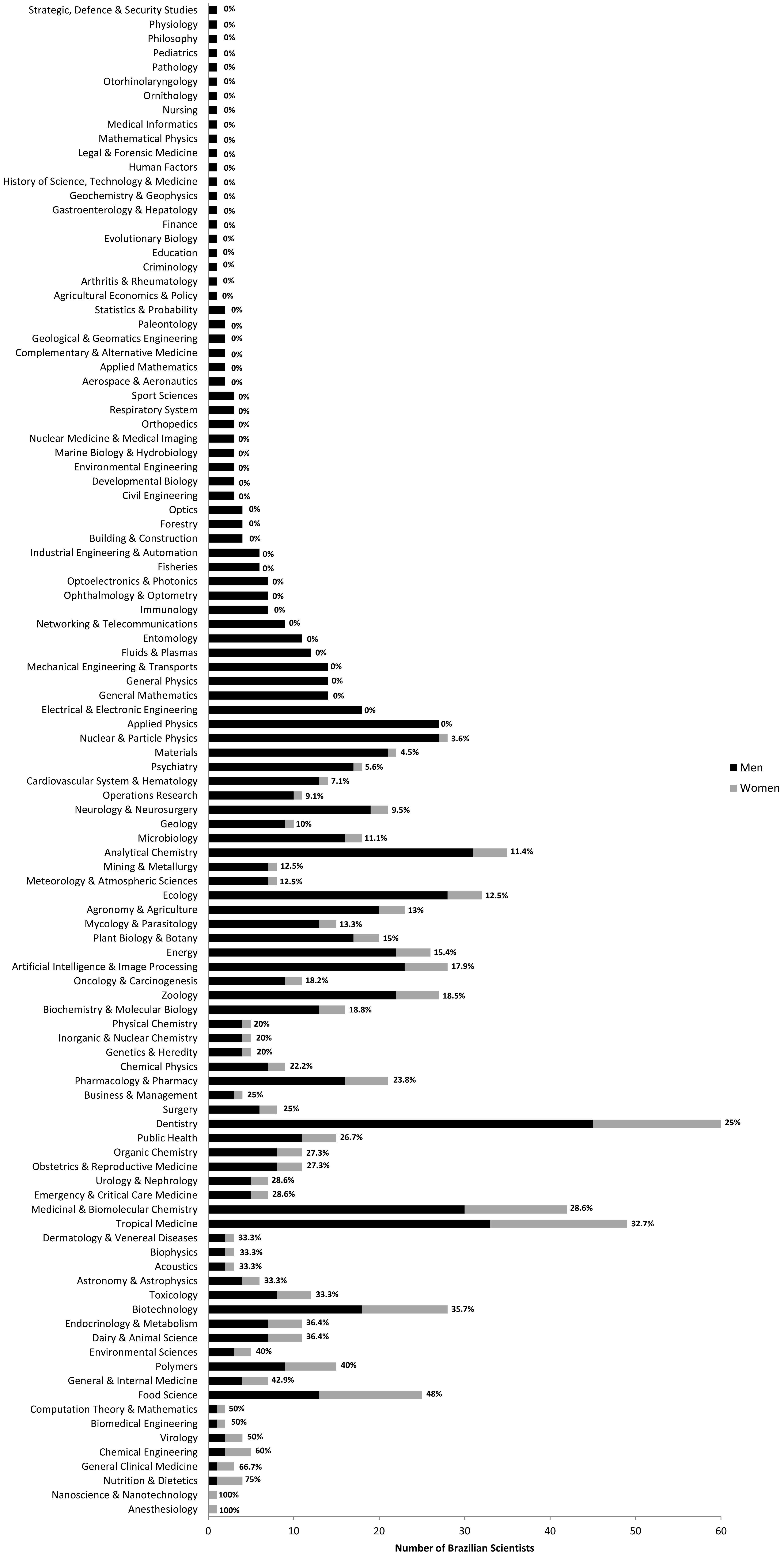
